# Does Long-Term Selection for Development Time Result in Canalization: A Test Using *Drosophila melanogaster*

**DOI:** 10.1101/553123

**Authors:** Shampa M. Ghosh, K. M. Satish, J. Mohan, Amitabh Joshi

**Author notes:** Correspondence: Dr. Shampa M. Ghosh Dr. Amitabh Joshi.

## Abstract

Canalization denotes the robustness of a trait against genetic or environmental perturbation. Plasticity, in contrast indicates the environmental sensitivity of a trait. Stabilizing selection is thought to increase canalization of a trait, whereas directional selection is often thought to lead to decanalization. However, the relationship between selection, canalization and plasticity remains largely unclear. Experimental evolution is a powerful approach for addressing fundamental questions in evolution. Here, we ask whether long-term directional selection for reduced pre-adult development time in *Drosophila melanogaster* results in the evolution of increased canalization for development time, the trait under primary selection. We additionally investigate whether pre-adult survivorship, a trait only secondarily under selection in this experimental regime, also evolves to become canalized. We examine canalization both in terms of stability of population means and of within population variability across two environmental axes. We used four large outbred populations of *D. melanogaster* selected for rapid pre-adult development and early reproduction for 295 generations, and four corresponding ancestral control populations that were not under conscious selection for development time or early reproduction. The selected populations had evolved 25% reduction in both development time and pre-adult survivorship at the time of this study. We studied development time and pre-adult survivorship of the selected populations and controls across various combinations ofrearing temperature and larval density. Development time in the selected populations had become more canalized than controls with regard to density, but not temperature. Canalization of development time across density appears to have evolved due to evolutionary changes in the lifehistory and physiology of the selected populations. Pre-adult survivorship, only a secondary correlate of fitness in the selected populations, did not show any clear trend in terms of canalization with regard to either density or temperature, and, overall variation in the trait was greater compared to development time within and across environments. Whether long-term directional selection canalizes or not, therefore, appears to be dependent in a complex way on specific interactions of trait, selection regime and environmental factor in the context of the ecology and physiology of the popualtions under study.

## Introduction

For organisms developing in the wild, environmental variation at various scales is ubiquitous; yet, the phenotypic expression of many traits shows a surprising degree of robustness (Félix and Barkoulas, 2015). This is thought to be a result of canalization (Waddington, 1942, 1961), the intrinsic ability of the developmental system to maintain its robustness in the face of environmental perturbations or underlying genetic variations (Waddington, 1942, 1961; Dworkin, 2005a). Buffering of the developmental process against environmental perturbation is referred to as environmental canalization (Waddington, 1956) and such perturbations could be macro or microenvironmental in nature (Falconer and Mackay, 1996; Debat and David, 2001). Macroenvironmental perturbations account for changes in factors like temperature, nutrition, social environment etc. during development. Microenvironmental perturbations, on the other hand, may arise due to small variations in the developmental environment over time and space, and on an even smaller scale because of stochastic fluctuations in the developmental process, also referred to as internal noise or developmental instability (Hall *et al.*, 2007; Morgante *et al.*, 2015). Developmental systems may show micro- or macroenvironmental canalization by maintaining their phenotypic constancy in the face of such perturbations. Genetic variation in development on the other hand, may arise in the form of new mutations, but there could be mechanisms that entrench the developmental process to generate robust phenotypes, masking the underlying genetic variation of a population (Waddington, 1956; Stearns and Kawecki, 1994; Wagner *et al.*, 1997; Rutherford and Lindquist, 1998; Sgrò *et al.*, 2010):this has been termed genetic canalization. By reducing expressed phenotypic variation, genetic canalization may help a population harbour hidden or cryptic genetic variation (Waddington, 1952, 1953; Bateman, 1959; Rendel, 1959; Gibson and Dworkin, 2004). Increase in trait variance of a population in a stressful environment is a common observation. This is thought to be due to breakdown of canalization in extreme environments (Dworkin, 2005a). Failure of the developmental buffering, or decanalization in extreme environments may reflect breakdown of genetic canalization and/or a lack of environmental canalization in the extreme environment. A trait that has reduced sensitivity to internal or external perturbations during development, and therefore shows a robust phenotypic expression despite perturbations or underlying variation, is said to be canalized.

Canalization or robustness as a phenomenon is intertwined with the related phenomenon of phenotypic plasticity. Phenotypic plasticity reflects the environmental sensitivity of a trait. Traditionally, plasticity referred to the ability of a genotype to give rise to different phenotypes in different environments (Woltereck, 1909). In more contemporary usage, a trait whose phenotypic expression varies in response to environmental variation is called a plastic trait, which could be simple (*e.g.* expression level of a protein), or complex (*e.g.* a life history trait) in nature. Plasticity and canalization of traits have often been thought of as opposite phenomena (Debat and David, 2001), as a trait which is highly canalized is, by definition, less plastic, and vice versa. However, a trait that is plastic with respect to a certain environmental variable could be canalized for another. Moreover, canalization of a trait expressed at one level of biological organization within an organism may require plasticity at an underlying or overlying level of organization (Proulx and Phillips, 2005; Dor and Jablonka, 2011; McDonald *et al.z*, 2018). For example, plasticity of a functional trait across enviroments may entail robustness of a higher-order trait like viability or fitness, leading to the view that both canalization and plasticity are best treated as aspects of the dependence of phenotype on the environment, being two sides of the same coin (de Visser *et al.*, 2003; Flatt, 2005). Studies on phenotypic plasticity have increased in the past couple of decades, and theis greater interest in finding the link between plasticity and evolution (West-Eberhard, 2003; Pigliucci, 2005; Fusco and Minelli, 2010; Moczek *et al.*, 2011; Paaby and Testa, 2018). The relationship between canalization and adaptive evolution, however, has remained largely unclear despite the ability of both processes to shape trait variation and hence influence each other (Debat and Rouzic, 2018; Groth *et al.*, 2018; Fossen *et al.*, 2018; Jiang *et al.*, 2018), In this paper we explore the link between adaptive evolution and canalization using a long-running selection experiment involving fruitflies, *Drosophila melanogaster*.

At a population level, both selection and canalization eliminate deviations from an optimal phenotype (Eshel and Matessi, 1998; Seigal and Bergman, 2002). Any mechanism that constrains the phenotype to be closer to the optimum is likely to be favoured by selection (Rendel, 1967; Stearns and Kawecki, 1994). Therefore, for a trait undergoing selection, the degree of its canalization is predicted to be positively correlated with its impact on fitness (Waddington, 1942; Schmalhausen, 1949). Although theoretical discussions related to this issue are quite old (Waddington, 1942; Schmalhausen, 1949), there is little empirical work done to test these predictions. Although studies of canalization are often focused on either morphological traits or gene expression levels (Dworkin, 2005b,c; Shaw *et al.*, 2014), life history traits are potentially helpful in exploring the evolutionary significance of canalization as they are directly connected to fitness (Prasad and Joshi, 2003; Garland and Rose, 2009). Stearns and Kawecki (1994) and Stearns *et al.* (1995) studied a suite of life-history traits like development time, body size, lifespan, and early and late fecundity in populations of *D. melanogaster* and found that the more important a trait is to fitness, the more strongly it is canalized against genetic and environmental perturbations. In our study, we focus on pre-adult development time and egg to adult survivorship in *D. melanogaster* using the approach of experimental evolution.

Experimental evolution is a powerful approach to address questions in evolutionary biology (Garland and Rose, 2009; Kawecki *et al.*, 2012). In this approach, replicate populations are allowed to evolve in the laboratory under specific selection pressures chosen by the experimenter, and the observed changes are studied over a number of generations. It allows the experimenter to observe adaptive evolution in real time, replicate the experimental set up to avoid interpreting stochastic effects as evolutionary outcomes, and draw statistically robust conclusions about evolutionary processes. In view of these, we used a long running experimental evolution study on *D. melanogaster* to explore the relationship between directional selection and canalization. Our study system involved four large outbred populations of fruitflies subjected to selection for rapid development and early reproduction, relative to ancestral controls, for close to 300 generations in the laboratory, and the four ancestral control populations from which the faster developing populations were selected (first described in Prasad *et al.*, 2000). The traits that we studied are pre-adult development time and survivorship from egg to adulthood.

Pre-adult development time had shown a strong response to selection with a 25% reduction in mean development time relative to ancestral controls after 100 generations of selection (Prasad *et al.*, 2001, Prasad and Joshi, 2003). After 245 generations, the magnitude of the selection response remained similar (Ghosh-Modak, 2009). Many other larval and adult traits exhibited correlated responses to selection for rapid development and early reproduction (Joshi *et al*., 2001; Prasad *et al.*, 2001, Prasad and Joshi, 2003; Shakarad *et al*., 2005; Ghosh-Modak, 2009; Ghosh-Modak *et al.*, 2009; Ghosh and Joshi, 2012; Dey *et al.*, 2016). In particular, pre-adult survivorship in the faster developing populations had reduced by about 25% compared to controls (Prasad *et al.* 2000; Ghosh-Modak 2009). In view of this, we examined the extent to which direct and correlated responses to directional selection experienced under stable environment are canalized, by investigating whether the selected populations exhibited canalization for development time and/or pre-adult survivorship across different rearing temperatures and egg densities, compared to their ancestral control populations.

Temperature and pre-adult density are known to affect both development time and survivorship in *Drosophila* (Prasad and Joshi, 2003). All of the eight (four selected, and four controls) populations used in this study have been maintained at a constant temperature and rearing density throughout the course of selection (Ghosh-Modak, 2009). For this particular study investigating canalization, these populations were reared in nine different environmental conditions, which comprised of three rearing temperatures crossed with three egg densities. One of the treatments corresponded to the standard maintenance conditions of the populations, and the remaining eight were novel environmental conditions to which the flies were not exposed before. We explore canalization and plasticity of the two traits (development time and pre-adult survivorship) in context of the selection pressures experienced by the populations used in the study, and discuss the possible causes for observed responses of the control and selected lines in specific environmental conditions.

## Materials and methods

### Experimental populations

We used eight laboratory populations of *D. melanogaster*: four replicate populations selected for rapid development and early reproduction, (FEJ _1-4_: **F**aster development, **E**arly reproduction, derived from **J**B, first described by Prasad *et al.* 2000), and their four matched ancestral control populations (JB _1-4_: **J**oshi **B**aseline, first described by Sheeba *et al.*, 1998). The four JB populations were descendants of a single wild-caught population of *D. melanogaster* — the IV population described by Ives (1970). They were first maintained in the laboratory for about 110 generations on a 14 day discrete generation cycle. Five populations (B_1-5_) were then derived from IV populations and reared in the laboratory under similar conditions (Rose and Charlesworth, 1981). After about 360 generations, a set of five populations were derived from B_1-5_ and christened UU_1-5_ (Uncrowded as larvae, Uncrowded as adults; described by Joshi and Mueller, 1996). The UU populations were maintained under similar conditions, but on a 21 day discrete generation cycle. After 170 generations of being maintained as UUs, JB_1-4_ populations were derived from UU populations (UU 1, 2, 3, 5, respectively) (Sheeba *et al.*, 1998). FEJ_1-4_ populations were later derived from JB_1-4_ and subjected to selection for rapid development and early reproduction, relative to their ancestral controls (Prasad *et al.* 2000).

All populations were maintained on a discrete generation cycle at ∼25°C, ∼90% relative humidity and constant light, on banana-jaggery food. In both JB and FEJ populations, larvae were reared in 8 dram glass vials (9.5 cm height, 2.4 cm inner diameter) with ~6 mL food at a density of 60–80 larvae per vial, whereas eclosed adults were collected into Plexiglas cages (25×20×15 cm^3^) with abundant food, at breeding population sizes of about 1500 flies. The JBs were maintained on a 3-week discrete generation cycle, and all eclosing adults were part of the breeding population. In JB populations, eggs were collected and dispensed in vials on the 21^st^ day after the egg collection for the previous generation, corresponding to day 11 of adult life. FEJs were maintained under conditions similar to the JBs, except that 120 vials, rather than 40 vials, containing approximately 60-80 eggs were collected per population, and the vials were monitored for eclosion every 2 h after pupae darkened. As soon as the first 20–25% flies in each vial (12-15 flies) had eclosed, they were transferred into fresh cages containing food plates. These constituted the breeding adults. After three days of adult life, eggs were collected from FEJ cages to start the next generation. Thus, the FEJs were under strong primary selection to complete egg-to-adult development fast, and under secondary selection to be relatively fecund on day 3 of adult life.

As each FEJ population was derived from one JB population, selected and control populations bearing identical numerical subscripts were more related to each other than to other populations in the same selection regime. Consequently, control and selected populations with identical subscripts were treated as constituting random blocks in the statistical analyses. At the time of this study, the FEJs had undergone 295 generations of selection, and showed considerable evolutionary reductions in

development time (∼25%), dry weight (∼50%), survivorship (~25%) and general level of activity (Prasad and Joshi 2003; Ghosh-Modak, 2009; Ghosh and Joshi, 2012).

### Collection of flies for assays

Prior to assays, all eight populations were reared under a common (control JB type) regime for one complete generation in order to ameliorate non-genetic parental effects. The eggs of these flies, hereafter referred to as standardized flies, were then used for the various assays.

### Development time assay

A fresh food plate was introduced into the cages and the standardized flies were allowed to lay eggs for one hour. This plate was then replaced by a second food plate and an egg collection window of one hour was provided. Eggs used for the assay were collected from the second round of egg lay and the first sets of plates were discarded. This was done to ensure that the eggs used for the assay were developmentally synchronized. Eggs were collected with a moistened paint-brush from the food-surface, counted under the microscope, and exact numbers of eggs for the assay were dispensed into vials with 6 mL of food at a density of 30, 70 or 300 eggs per vial and incubated at three different temperatures namely 18°C, 25°C and 28°C. 25°C and 70 eggs per 6 mL of food represent the normal maintenance conditions for FEJ and JB populations. The egg density of 30 eggs per vial was considered in order to study development at density substantially lower than what the populations normally experienced. 300 eggs in 6 mL, on the other hand, represents substantial larval crowding for these *Drosophila* populations, and the choice of a density that could be somewhat stressful was deliberate as canalization of development or the lack thereof could be explored.

Eight vials were set up for each combination of temperature, density, selection regime and replicate block (total 72 treatments). In all, 576 vials (2 selection regimes × 4 replicate blocks × 3 temperatures × 3 densities × 8 vials) were thus set up for the experiment. The vials were monitored for the first eclosion and, thereafter, checked regularly at 4 h intervals and the number of eclosing flies was recorded. The observations were continued till no new fly eclosed for two consecutive days. The development time in hours was measured by subtracting the time of egg-lay from the time of eclosion. In 4 out of the 72 treatments (2 selection regimes × 4 replicate populations × 3 temperatures × 3 densities), one vial was not viable, possibly due to handling error. For these sets, only data for 7 vials were available. To maintain a balanced design for the statistical analyses, for all 72 treatments the data from one vial, chosen at random, were excluded from the analyses. Hence, data from a total of 504 vials were considered for the final analyses.

### Egg to adult survivorship assay

From the vials used for the development time assay, the total number of eclosed flies were counted, and the number was divided by the number of eggs (30, 70, 300) for the respective vial to obtain the egg to adult survivorship for any given vial. Similar to development time, survivorship data from a total of 504 vials were considered for analyses.

### Outlier removal

Being large and outbred populations, there is considerable variation for development time, especially within the JB, and once eclosion starts for a given population, it roughly continues for 24 to 48 h. A temperature of 25°C and density of ~70 eggs per vial represent the standard developmental condition of the JB and FEJ populations. Under such conditions, FEJ flies take 6 to 7 days to complete their development from egg to eclosion, whereas JB flies take 9 to 10 days to complete pre-adult development. For the development time data analysis, flies eclosing after 100 hours (over 4 days) from the first eclosion for any given set (combination of particular selection regime, temperature, density and replicate population) were not included in the development time data analysis for 30 and 70 egg densities. For 300 egg density, flies eclosing after 200 hours (over 8 days) from the first eclosion for any given set were not included in the development time analysis. The criteria for defining outliers, typically few in number and representing potentially pathological individual variants, was decided upon before the experiment, based on past experience of development time distributions in these populations. Ultimately, outlier removal had to be done for only 6 of the 72 treatment combinations and essentially implied the exclusion of one or two extremely late developing flies as outliers in any given vial. For survivorship data, all eclosed flies, including the outliers defined as above, were considered for the analysis as inclusion of these flies did not greatly increase the mean or variance for survivorship.

### Statistical analyses

For both pre-adult development time and survivorship, we examined the mean trait value, and variation in trait values, in each of the eight populations in all nine combinations of rearing density and temperature. We quantified plasticity or macroenvironmental sensitivity of a trait as the relative change in its value from one assay environment to another. For a given selection regime, less plasticity of mean trait value across assay environments would indicate greater macroenvironmental canalization for a trait.

For comparing trait variation across selection regimes and environments, we used the *coefficient of variation* instead of *variance*. The coefficient of variation or CV is a standardised, dimensionless quantity measured by dividing the sample standard deviation by the sample mean (often expressed as a percentage and hence, multiplied by 100). Coefficients of variation is used specifically when comparing the variation of two *populations* independent of the magnitude of their means, or alternatively, while comparing the variations in two *traits* irrespective of their means (Sokal and Rohlf, 1998). Prior data suggest both development time and survivorship of the faster developing populations used in our study are ~25% less than that of the ancestral controls, and development time is also less variable in selected populations, compared to controls (Prasad *et al.* 2000; Prasad *et al.*, 2001; Ghosh-Modak, 2009). Hence, we used CV as a standardised measure of varaibility for both development time and survivorship in all eight study populations.

Mixed-model analyses of variance (ANOVA) were performed on all data, using a completely randomized block design in which selection regime, temperature and density were treated as fixed factors, crossed among themselves, and with random blocks, representing both ancestral lineage and coincident handling of one matched pair of replicate selected and control populations during assays. For analyzing across-environment variation in mean trait values, we used vial mean values for development time and vial values for pre-adult survivorship as the input data for the analysis. For comparing development time variation across individuals, the values of CV across individuals within each vial were used as input data for ANOVA. In addition, the across vial variability in mean development time for all combinations of block, selection regime, temperature and density was also subjected to ANOVA. For survivorship, only vial means could be scored given the nature of the trait, and CV for each vial could not be calculated. Hence, only the across vial variability was used as a measure of survivorship variability for any population and treatment. All analyses were implemented using the software JMP (SAS institute) version 14. For post hoc comparisons, Tukey’s HSD test was performed using JMP. All significant results in the post hoc comparisons indicate a *p-*level less than 0.05. Since the basic consequential units of analysis in this design are mean values of the replicate populations (e.g. MS selection regime will be tested over MS block × selection regime interaction, etc), normality can be safely assumed, even though traits like development time are not distributed normally across individuals.

## Development time data analysis

### Mean development time (trait mean)

*Mean* development time was calculated for each vial. A four way mixed model ANOVA was performed on vial means as within-cell replicate values, with selection, temperature, and density being treated as fixed factors, and block An effect of treatment combination (temperature and density) would then be taken as reflecting macroenvironmental sensitivity of the trait.

### Trait variation across individuals (within vials)

Trait variation observed in different combinations of temperature and density in the eight populations was also subjected to four-way mixed model ANOVA. Variation among individuals within a vial can reflect genetic variation among individuals, developmental instability which is hard to distinguish from variation due to possible ultra-microenvironmental heterogeneity within a vial, in addition to stochastic variation in trait measurement. Partitioning the within vial variation in phenotype among individuals into components caused by these four factors cannot be accomplished. However, in the context of our experimental design, if different combinations of the macroenvironmental factors, temperature and density, affect the extent of within vial variation in phenotype among individuals, it can be cautiously interpreted as a change in developmental instability and/or ultra-microenvironmental sensitivity, given that the macroenvironmental change is unlikely to affect genetic variation or stochastic error. Incidentally, the one time the FEJ and JB populations were examined for developmental instability for any trait, no significant differences were observed (Shakarad *et al*. 2001). In that study, the trait examined was fluctuating asymmetry of sternopleural bristle number. For the current study, two different measures of trait variation within a population in any specific treatment combination were used. The first one was the standard deviation of development time across individuals, within a vial. The second was CV or coefficient of variation for the trait within each vial. Standard deviations of individual development time were calculated for each vial and divided by the respective vial means for development time to obtain replicate measures of CV, expressed as a percentage, for the trait.

### Trait variation across vials

Across vial variation for development time reflects the effects of sampling, since there is within vial variation among individuals, overlaid by effects of microenvironmental heterogeneity, if any. There is evidence for such among vial microenvironmentally induced variation in life-history traits in *Drosophila* populations such as these in our laboratory (Dey *et al*., 2006). Again, while it would not be possible to tease apart these two effects within a population, the effects of combinations of the macroenvironmental factors, temperature and density, on among vial variation can be cautiously assumed to be operating primarily through an effect on the microenvironmental sensitivity, since macroenvironment is unlikely to affect sampling variation. Again, two measures of among vial variation were used, and tested via four-way mixed-model ANOVA: standard deviation of vial means for a particular treatment (selection regimes × replicate populations × temperature × density) and CV (standard deviation of vial means divided by grand mean for the treatment), multiplied by 100.

### Survivorship data analysis

Data on pre-adult survivorship were treated exactly as described above for development time, except that mean pre-adult survivorship for each population was calculated by averaging the per vial pre-adult survivorship, followed by an arcsin square-root transformation. Since each vial yielded only one pre-adult survivorship value, only among vial variation was examined. The inference of macroenvironmental and microenvironmental sensitivity changes in populations subjected to different combinations of temperature and density was exactly the same as for development time.

## Results

### Egg to adult development time

#### Mean egg to adult development time across macroenvironments

Mean development time of FEJ is significantly less (~75%) than that of JB (F_1,3_ = 5372.666; p < 0.0001) across all combinations of temperature and density (Figure 1, Table 1). Interaction between selection regime and density is significant (F_2,6_ = 81.8427; p < 0.0001) (Table 1). At 30 and 70 eggs per vial, the FEJ flies develop 25% faster than that of JB, across rearing temperature (Figure 1). At a higher rearing density of 300 eggs per vial though, FEJ populations develop 30% faster than JB, due to JB development getting slower at 300. The post hoc comparisons show that for any given temperature, mean development time in FEJ does not different significantly across density. In contrast, mean development time in JB increases by ~10% from 70 to 300 egg density, averaged across temperatures, and this change is significant (p < 0.05). Mean development time in JB is not significantly different between 30 and 70 egg density. Hence, a culture density of 300 eggs per vial delays JB development significantly but not FEJ. Thus, from a density of 70 to 30 eggs per vial, the *selection response* is canalized, but not when density increases from 70 to 300 eggs per vial.

**Table 1:** Results of ANOVA on mean development time. Only fixed factor effects could be tested for significance.

| **Effect** | ***df*** | ***F*** | ***P*** |
| --- | --- | --- | --- |
| Selection | 1 | 5372.666 | <.0001* |
| Density | 2 | 219.1276 | <.0001* |
| Selection × density | 2 | 81.8427 | <.0001* |
| Temperature | 2 | 2215.963 | <.0001* |
| Selection × temperature | 2 | 1358.630 | <.0001* |
| Density × temperature | 4 | 11.3361 | 0.0005* |
| Selection × density × temperature | 4 | 14.0372 | 0.0002* |

**Figure 1:**
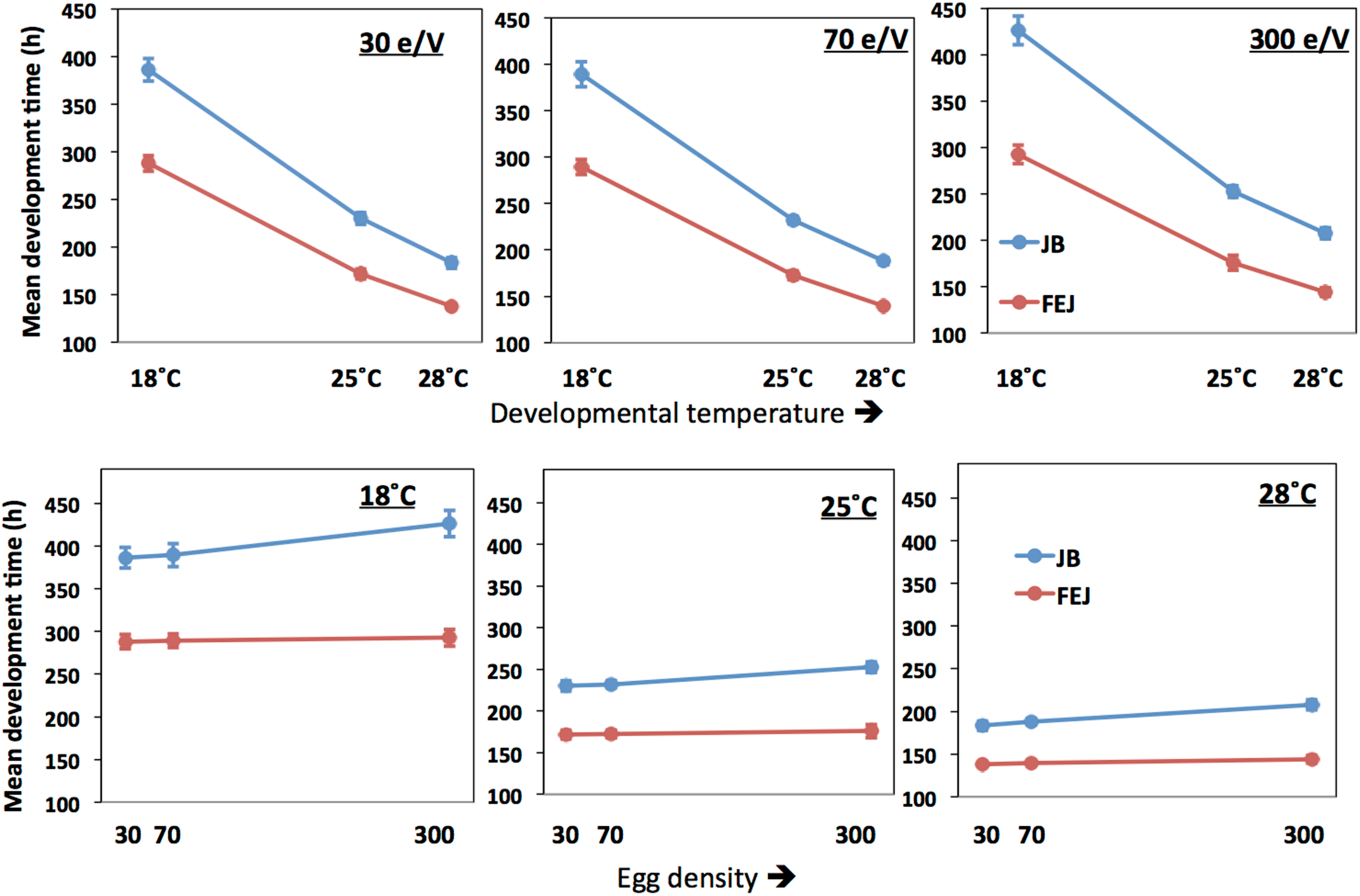
Mean egg to adult development time of FEJ and JB populations. The upper panel shows mean trait value plotted across temperature for each treatment density (e/V = eggs per vial). The lower panel shows mean trait value plotted across egg density for each treatment temperature. The error bars show 95% confidence intervals calculated from the variation among replicate populations in each combination of selection regime, temperature and density.

In both FEJ and JB, mean development time changes significantly across temperature for any given density but both JB and FEJ show similar changes across temperature. Mean development time in JB and FEJ increases by 68% and 67% respectively from 25°C to 18°C, averaged across all densities (Figure 1). Mean development time in both JB and FEJ reduces by 19% from 25°C to 28°C, averaged across all densities (Figure 1). Thus, the thermal plasticity of mean development time is consistent across densities and selection regimes, resulting in the selection response being canalized across the tested temperature range, from 18°C to 28°C.. However, there is significant interaction between selection and temperature (F_2,6_ = 1358.630; p < 0.0001), as JB development time shows greater absolute change across temperature (Figure 1, Table 1).

To summarize, overall, mean development time is more plastic along the temperature axis than the density axis (Figure 1). FEJ development is faster than that of the JB, and relatively more canalized than that of JB across rearing densities. Mean development time, although being highly plastic across temperature, responds similarly to changes in temperature in both the FEJ and JB populations. Thus, though development time as a trait is less canalized (more plastic) with respect to temperature as compared to density, the *selection response* of development time is actually more canalized with regard to temperature than density.

### Across vial (microenvironments) variation for egg to adult development time

The standard deviation of (vial mean) development time across vials is significantly lower in FEJ than in JB (F_1,3_ = 15.5816; p = 0.029), and is also significantly affected by temperature, being higher at 18°C compared to the higher temperatures in both the JB and FEJ populations (F_2,6_ = 25.98; p = 0.001) (Table 2). However, both these main effects in the ANOVA appear to be due to the standard deviation scaling with the mean. When we examined the CV of development time across vials, there was no significant effect of any factor or interaction in the ANOVA (Table 3), and in both FEJ and JB populations, the CV of development time across vials was about 1, regardless of temperature or rearing density (Figure 2). Thus, the sensitivity of development time to microenvironmental variation does not seem to be affected either by the two macroenvironmental variables (temperature and density), or by selection history.

**Table 2:** Results of ANOVA on standard deviation of development time across vials. Only fixed factor effects could be tested for significance.

| **Effect** | ***df*** | ***F*** | ***P*** |
| --- | --- | --- | --- |
| Selection | 1 | 15.5816 | 0.0290* |
| Density | 2 | 4.2555 | 0.0707 |
| Selection × density | 2 | 2.9859 | 0.1259 |
| Temperature | 2 | 25.9890 | 0.0011* |
| Selection × temperature | 2 | 1.7682 | 0.2491 |
| Density × temperature | 4 | 1.8152 | 0.1908 |
| Selection × density × temperature | 4 | 0.8867 | 0.5009 |

**Table 3:**
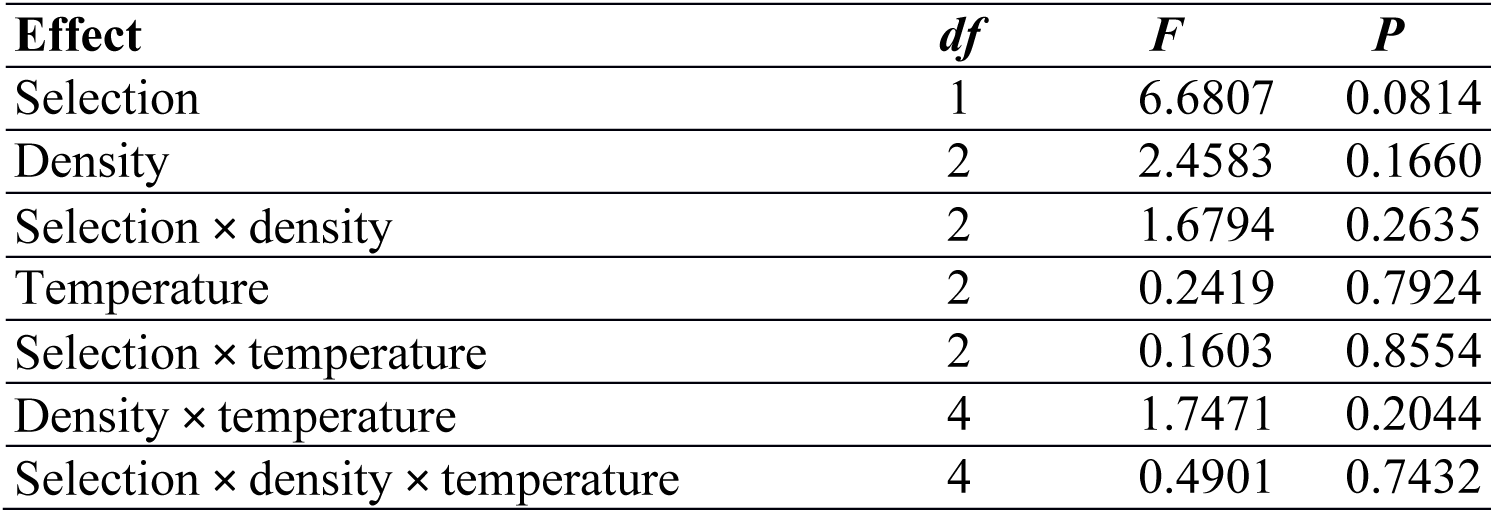
Results of ANOVA on CV of development time across vials. Only fixed factor effects could be tested for significance.

**Figure 2:**
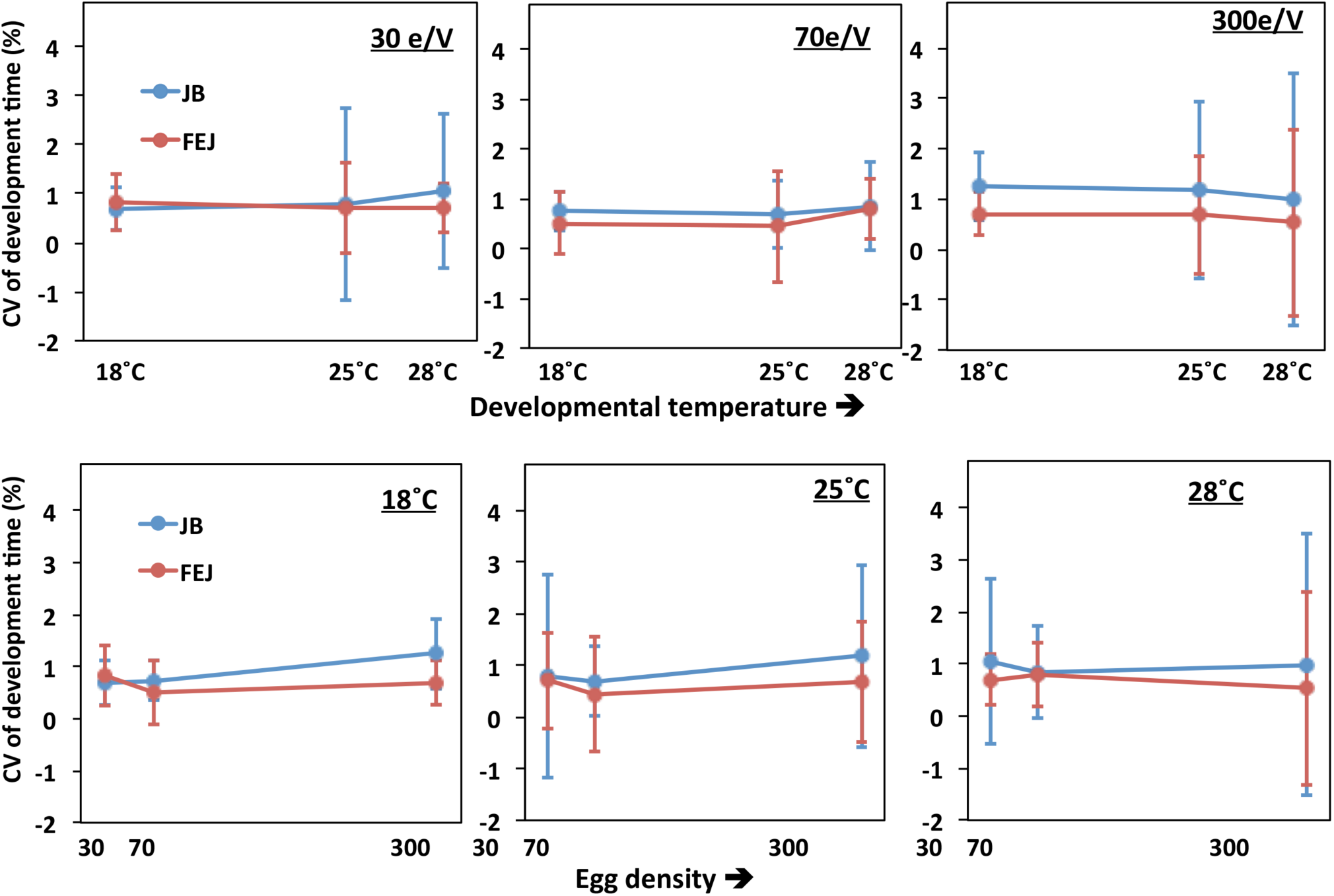
Coefficient of variation (CV) of mean egg to adult development time of FEJ and JB populations acrossvials. The upper panel shows CV of the trait plotted across temperature for each treatment density (e/V = eggs per vial). The lower panel shows CV of the trait plotted across egg density for each treatment temperature. The error bars show 95% confidence interval calculated for the replicate population (block) means of CV.

### Across individual variation (within vial CV) for egg to adult development time

Overall, the among individual within vial CV in development is significantly less in FEJ compared to JB (F_1,3_ = 16.7120; p = 0.0265) (Table 4). The ANOVA also revelaed significant main effects of density (F_2,6_ = 1397.088; p < 0.0001) and temperature (F_2,6_ = 25.883; p = 0.0011), with CV tending to increase with increases in both temperature and density, though the temperature effect was less consistent than that of density and seen prominently only at the highest rearing density of 300 eggs per vial (Figure 3). These patterns are reflected in significant interactions between selection and density (F_2,6_ = 24.9087; p = 0.0012), and density and temperature (F_4,12_= 11.882; p = 0.0004). Among individual CV for development time at all three temperatures is significantly lower in FEJ compared to JB only when reared at a density of 300 eggs per vial (p < 0.05), but not at 30 or 70 eggs per vial (Figure 3). CV of JB development time increases by a staggering 138%, on average, from 70 to 300 egg density, and the changes are similar at all temperatures. CV of FEJ development time, on the other hand, increases by only by 22% from 70 to 300 egg density at 18°C, which is not significant. At 25°C and 28°C, changes in the CV of FEJ development time are 98% and 78% respectively when the density changes from 70 to 300, which are significant (Figure 3). Hence, the CV in FEJ development time is similar to that in JB at lower developmental densities, but at 300 eggs per vial, the CV in FEJ development time is significantly less than that in JBs as the FEJs undergo a much smaller increase in CV between 70 and 30 eggs per vial (Figure 3). CV for development time in both FEJ and JB is siginificantly greater at 28°C than 18°C only when reared at 300 egg density (p < 0.05). Overall, both FEJ and JB populations are similarly affected by temperature (no significant selection by temperature interaction, Table 4), and the temperature effect is greatly enhanced at the highest density of 300 eggs per vial (Figure 3; significant density by temperature interaction, Table 4).

**Table 4:** Results of ANOVA on CV (coefficient of variation) of development time across individuals, within vials. Only fixed factor effects could be tested for significance.

| **Effect** | **df** | **F** | **P** |
| --- | --- | --- | --- |
| Selection | 1 | 16.7120 | 0.0265* |
| Density | 2 | 397.0883 | <.0001* |
| Selection × density | 2 | 24.9087 | 0.0012* |
| Temperature | 2 | 25.8835 | 0.0011* |
| Selection × temperature | 2 | 0.7155 | 0.5264 |
| Density × temperature | 4 | 11.8821 | 0.0004* |
| Selection × density × temperature | 4 | 0.3931 | 0.8097 |

**Table 5:** Results of ANOVA for mean survivorship. Only fixed factor effects could be tested for significance.

| **Effect** | **df** | **F** | **P** |
| --- | --- | --- | --- |
| Selection | 1 | 133.7525 | 0.0014* |
| Density | 2 | 135.7491 | <.0001* |
| Selection × density | 2 | 6.6117 | 0.0304* |
| Temperature | 2 | 0.9151 | 0.4499 |
| Selection × temperature | 2 | 5.6709 | 0.0414* |
| Density × temperature | 4 | 1.3094 | 0.3213 |
| Selection × density × temperature | 4 | 0.7017 | 0.6056 |

**Figure 3:**
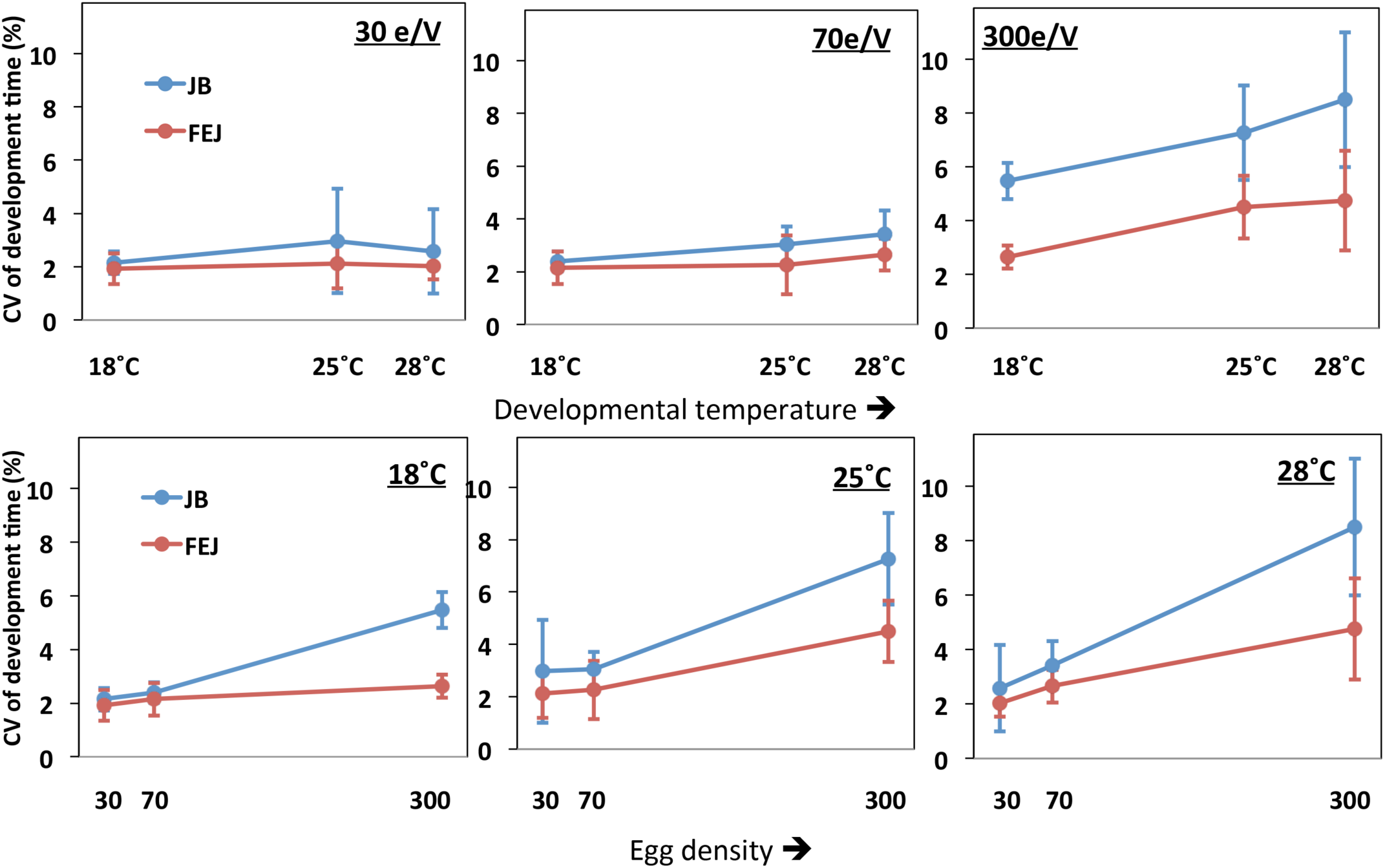
Coefficient of variation (CV) of egg to adult development time of FEJ and JB populations across individuals within a vial. The upper panel shows CV of the trait plotted across temperature for each treatment density (e/V = eggs per vial). The lower panel shows CV of the trait plotted across egg density for each treatment temperature. The error bars show 95% confidence intervals calculated from the variation among replicate populations in each combination of selection regime, temperature and density.

### Egg to adult survivorship Mean egg to adult survivorship

The pattern of macroenvironmental effects on mean egg to adult survivorship in the FEJ and JB is not as consistent as in the case of development time. Overall, mean survivorship in FEJ was less than in JB (F_1,3_ = 133.752; p = 0.0014), as also noted in earlier studies (Prasad *et al.*, 2000; Ghosh and Joshi, 2012). There was also a significant main effect of density (F_2,6_ = 135.749; p < 0.0001), with survivorship dropping markedly at 300 eggs per vial for both selection regimes, compared to that at 30 or 70 eggs per vial (Figure 4). Moreover, selection regime also interacted significantly with both temperature (F_2,6_ = 5.6709; p = 0.0414) and density (F_2,6_ = 6.6117; p = 0.0304), with mean survivorship in FEJ being significantly less than that of JB at 18°C and 28°C, but not at 25°C, although the difference is substantial even at 25°C, except at a density of 300 eggs per vial (Figure 4).

**Figure 4:**
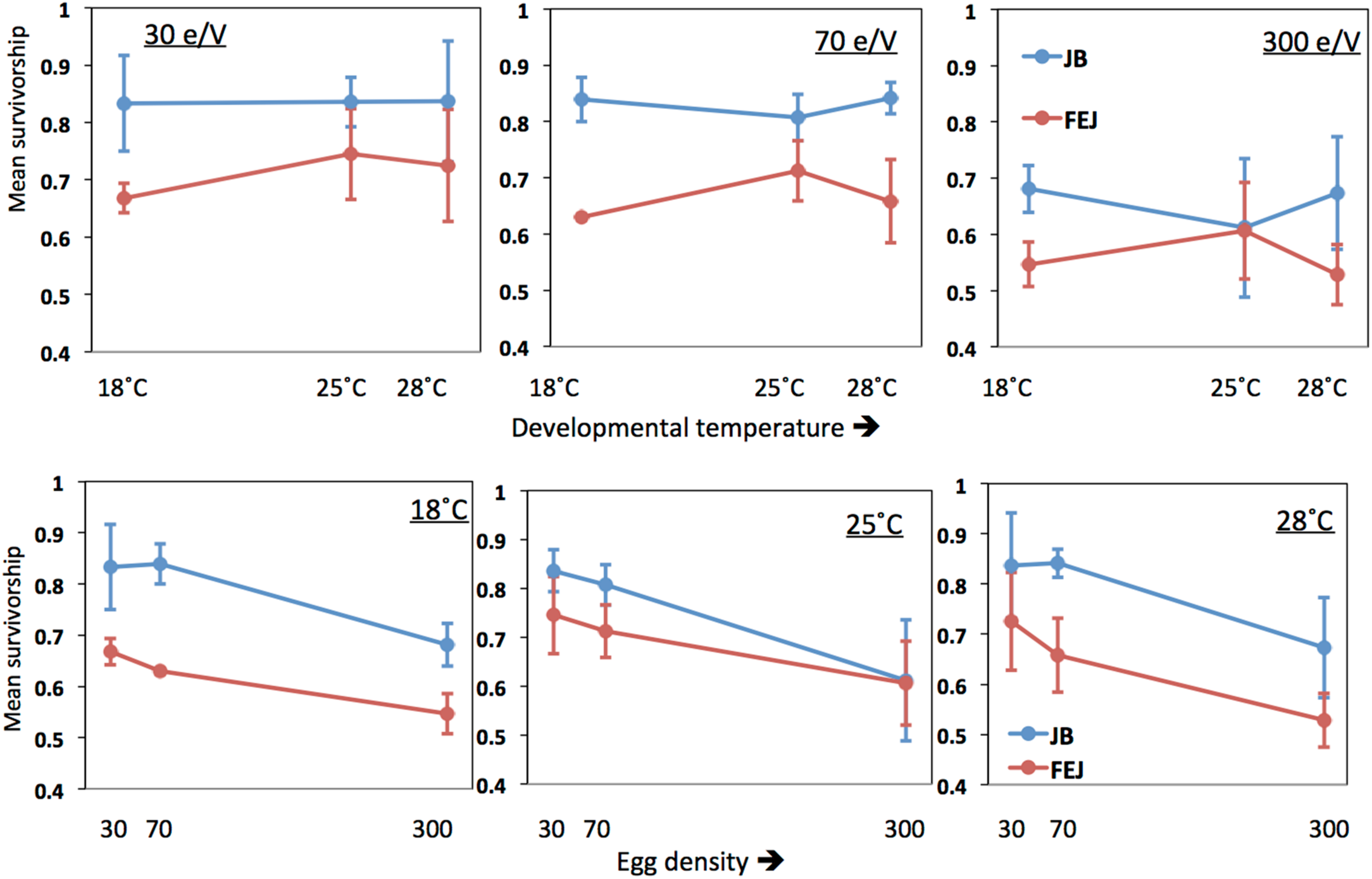
Mean egg to adult survivorship of FEJ and JB populations. The upper panel shows mean trait value plotted across temperature for each treatment density (e/V = eggs per vial). The lower panel shows mean trait value plotted across egg density for each treatment temperature. The error bars show 95% confidence intervals calculated from the variation among replicate populations in each combination of selection regime, temperature and density.

Mean survivorship in both JB and FEJ is significantly reduced at a density of 300 eggs per vial compared to either of the two lower densities (Figure 4). However, at 25°C and 18°C, FEJ undergo a smaller reduction in mean survivorship (14%, 13%) compared to JB (24%, 19%) when density is increased from 70 to 300 eggs per vial. At 28°C, the reduction in survivorship from 70 to 300 density is similar (20%) in FEJ and JB. Overall, mean survivorship seems to be more affected by changes in density than temperature.

### Across vial variation (standard deviation and CV) for egg to adult survivorship

CV of survivorship across vials is consistently and substantially higher than that of development time for both JB and FEJ populations (Figure 5 *vs*. Figure 2; Figure 6), suggesting that survivorship is more sensitive than development time to microenvironmental variation across vials. For both standard deviation and CV, the only significant ANOVA effects were those of density and the selection by density interaction (Tables 6,7). Averaged across selection regimes, the CV of survivorship tended to decrease with increasing density, though the bigger drop was from 30 to 70 eggs per vial (Figure 5). However, in the FEJ, CV of survivorship tended to drop substantially from 30 to 70 to 300 eggs per vial, whereas in the JB, CV of survivorship was the least at 70 eggs per vial (Figure 5).

**Table 6:**
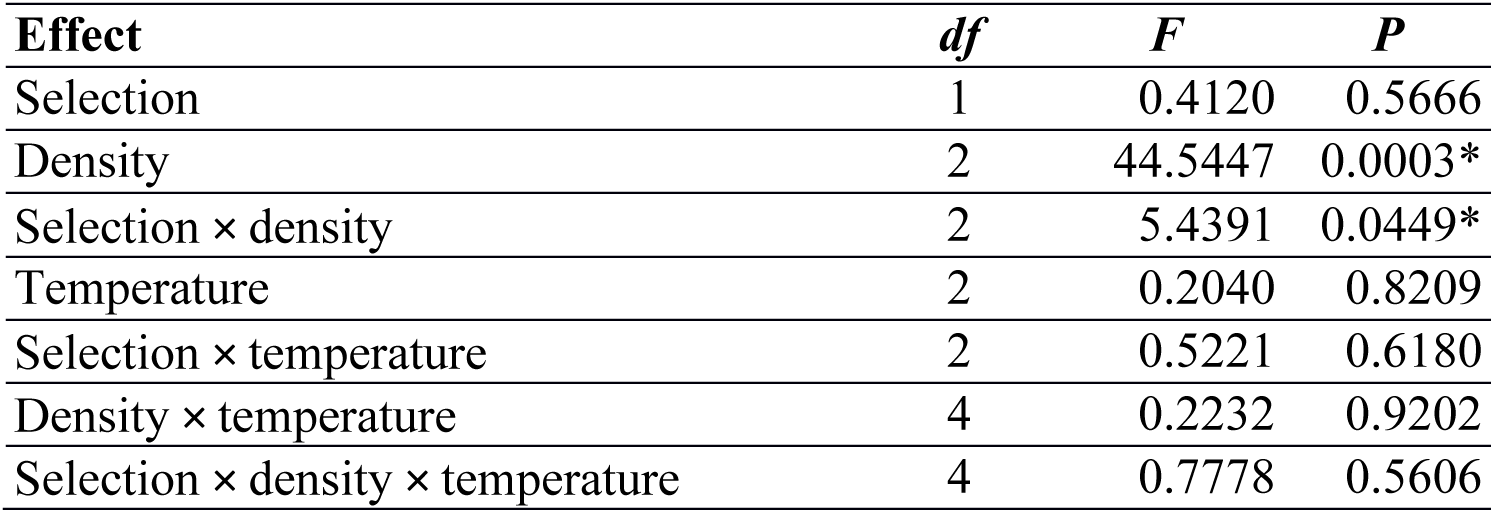
Results of ANOVA for standard deviation of survivorship across vials. Only fixed factor effects could be tested for significance.

**Table 7:** Results of ANOVA for CV of survivorship across vials. Only fixed factor effects could be tested for significance.

| **Effect** | **df** | **F** | **P** |
| --- | --- | --- | --- |
| Selection | 1 | 1.5481 | 0.3018 |
| Density | 2 | 16.3390 | 0.0037* |
| Selection × density | 2 | 7.0492 | 0.0266* |
| Temperature | 2 | 0.0853 | 0.9193 |
| Selection × temperature | 2 | 1.1776 | 0.3703 |
| Density × temperature | 4 | 0.0821 | 0.9864 |
| Selection × density × temperature | 4 | 0.9375 | 0.4750 |

**Figure 5:**
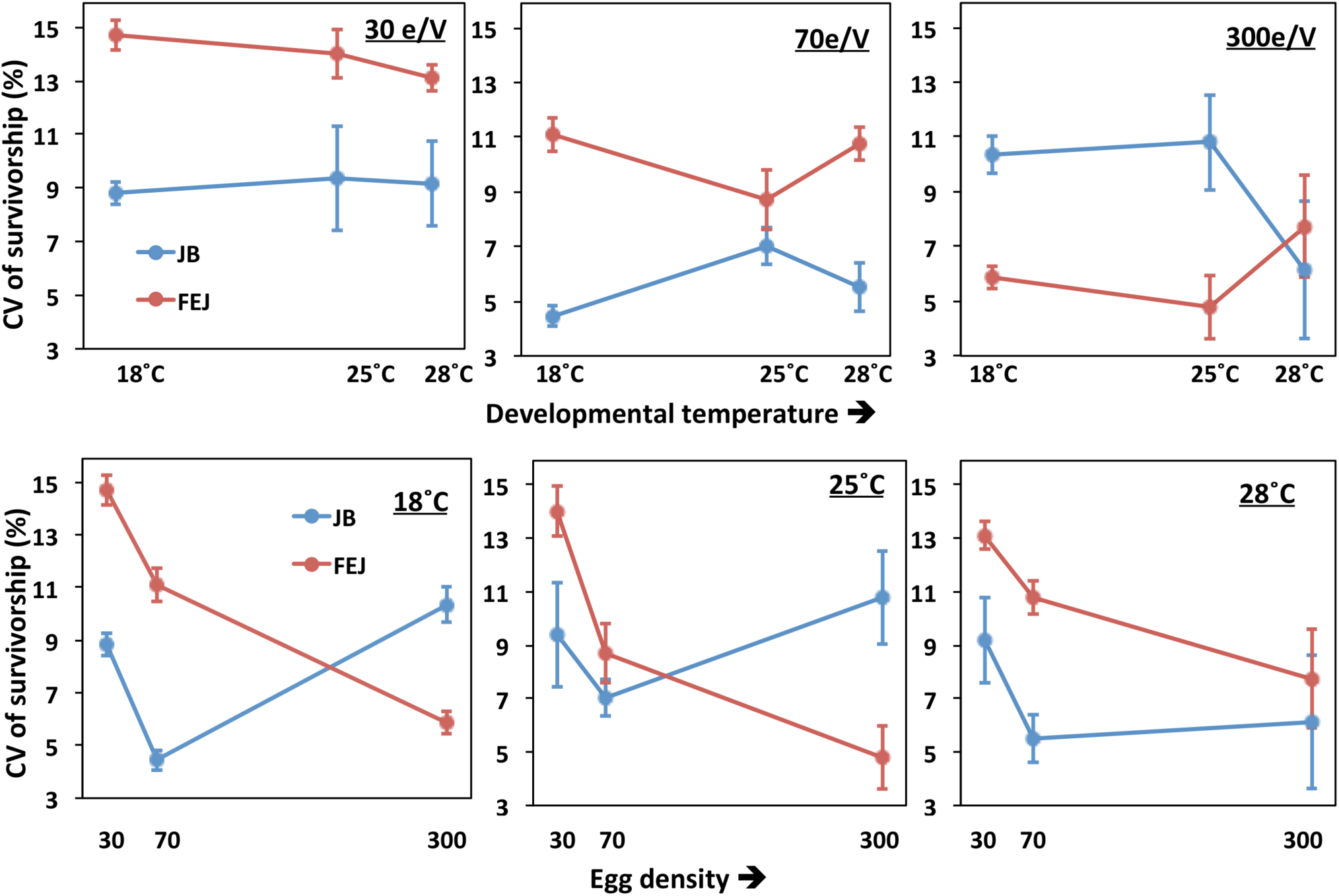
Coefficient of variation (CV) of egg to adult survivorship of FEJ and JB populations across vials. The upper panel shows CV of the trait plotted across temperature for each treatment density (e/V = eggs per vial). The lower panel shows CV of the trait plotted across egg density for each treatment temperature. The error bars show 95% confidence intervals calculated from the variation among replicate populations in each combination of selection regime, temperature and density.

**Figure 6:**
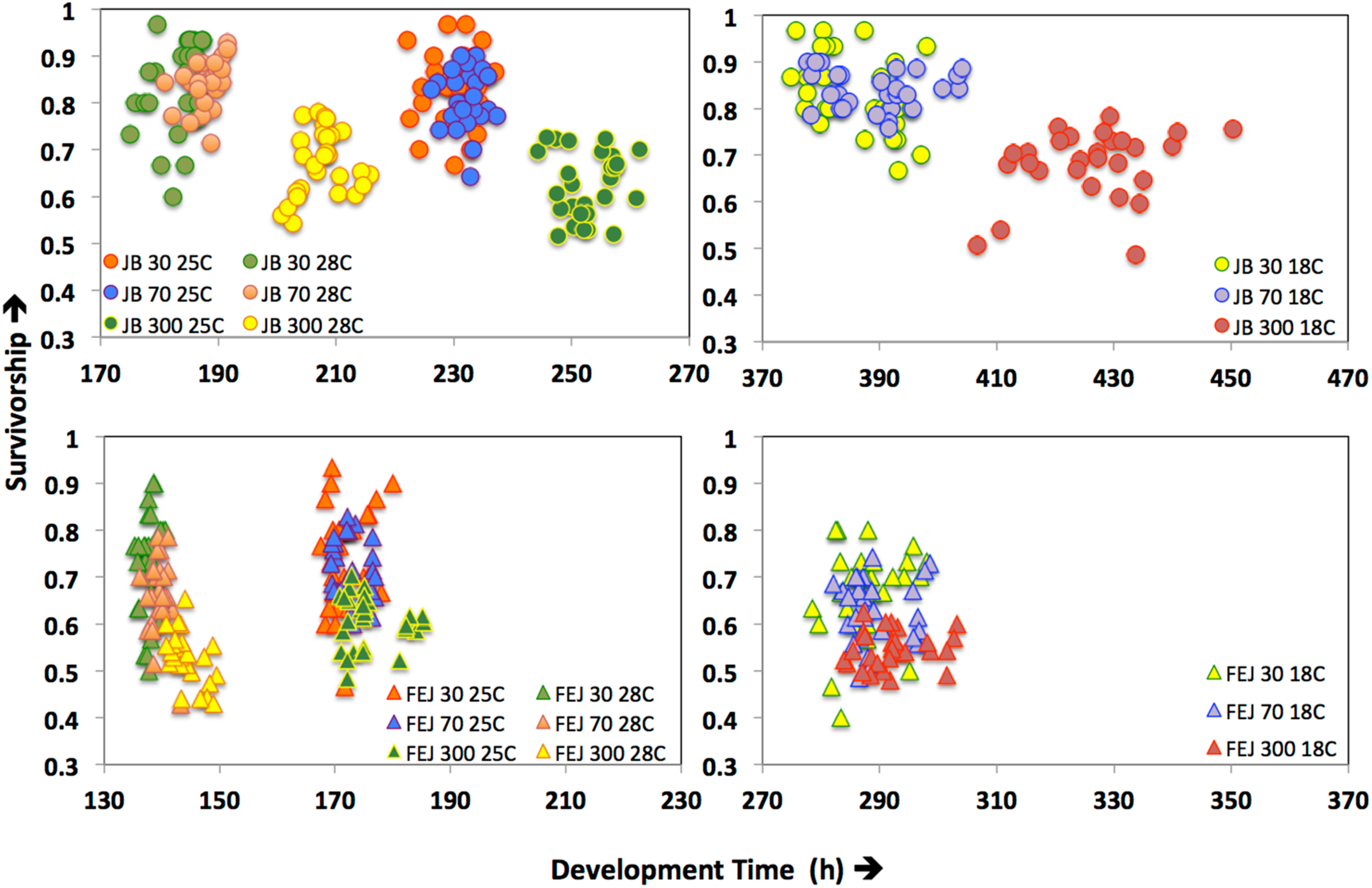
Plot of vial values of survivorship (y-axis) *vs.* vial mean values of development time (x-axis) for FEJ and JB populations across all treatments. Each circle represents value or mean obtained from one vial.

Thus, CV of egg to adult survivorship across vials is affected more strongly by changes in density than temperature (Figure 6), with different patterns in FEJ and JB. Unlike in the JB, CV of survivorship in FEJ is much higher at the lowest density of 30 eggs per vial (Figures 5 and 6). Survivorship from egg to adult is known to be reduced at very low densities in *Drosophila* due to the food medium not getting softened by the feeding activity of enough larvae, and also reduced suppression of fungal growth by larvae (<u>Sang, 1956</u>; Ashburner, 1989). It is possible that egg to adult survivorship in the FEJ is more sensitive to across vial microenvironment variation because the FEJ larvae, being much slower feeders than JB larvae (Prasad *et al*. 2001), exhibit an Allee effect even at a density of 30 eggs per vial, whereas that density is sufficiently high for JB larvae not to be exhibiting an Alle effect.

## Discussion

The structure of our experiment permits us to make several pair-wise comparisons of the effects of long-term directional selection on the sensitivity of fitness-related traits to micro- and macroenvironmental effects. Before drawing these contrasts, and discussing some of the implications for our understanding of the relationship between selection and canalization, we first briefly summarize the patterns seen in our results when we examined egg to adult development time and egg to adult survivorship in the selected populations (FEJ) and their ancestral controls (JB). It should also be noted that though both development time and survival to adulthood are fitness components, it is development time that has the higher correlation with fitness in the FEJ selection regime, with egg to adult survivorship being a secondary fitness correlate.

In terms of the macroenvironmental effects of rearing temperature and density on mean development time in the FEJ and JB, temperature had greater effects than density on mean development time (Figure 1), indicating that the thermal plasticity of this trait is greater than its plasticity to density. Temperature and development time are inversely related in ectotherms (van der Have and de Jong, 1996; Angilletta *et al.*, 2004; Ghosh *et al.*, 2013) and this relationship is well known in *Drosophila* (Gebhardt and Stearns, 1998; Gibert *et al.*, 2004). Likewise, in our study, mean development time increased at 18°C, and became shorter at 28°C, compared to 25°C, the standard rearing temperature of both FEJ and JB (Figure 1). Although the JB development time underwent greater absolute change with changing temperature, the relative changes in development time compared to the standard, *i.e.* 25°C, were same in both JB and FEJ. Thus, despite a large evolutionary reduction of development time in FEJ populations, thermal (macroenvironmental) plasticity of development time has not evolved in FEJ populations.

Like temperature, larval density is a major macroenvironmental variable affecting egg to adult development time in *Drosophila*, but in a more complex manner. At very low larval densities (10 or so larvae per vial), *Drosophila* cultures show an Allee effect, which signifies the beneficial effects of larger numbers of larvae to work up and soften the food medium and control the growth of fungi in absence of which development and survival may be affected for the existing larvae (Sang, 1956; Ashburner, 1989; A. Joshi, *pers*. *obs*.). Beyond a moderate density (50-100 larvae per vial), increasing larval density tends to increase mean development time, largely by stretching out the right tail of the development time distribution (Borash *et al.*, 1988; Sarangi, 2018) Our results show that mean development time of FEJ populations remains more uniform across rearing densities compared to the JBs, the difference being especially visible at the highest density of 300 eggs per vial, and at the lowest temperature of 18°C (Figure 1). As the egg density rises from 70 to 300 eggs per vial, relative increase in mean development time is 4 times higher in JB than in FEJ, indicating macroenvironmental canalization of development time with regard to larval density in the FEJ, relative to JB controls. We believe the reason for this canalization lies in the reduction of the third instar in FEJ larvae (Prasad *et al*. 2000). FEJ larvae cease feeding and start wandering in search for pupariation sites soon after reaching their minimum critical size, whereas JB larvae continue to feed for a long time after critical minimum size is attained; this pattern is mirrored in very low levels early in the third instar of FEJ of the transcript of the *Dnpf* (*Drosophila* neuropeptide F) gene, reduction in expression of which triggers the shift from feeding to wandering, typically late in the third instar (Satish, 2010). Moreover, as a consequence, FEJ larvae are much smaller than JB larvae at pupariation, and consume less food during the larval stage, resulting in increased carrying capacity (Joshi *et al.*, 2001). Thus, the spreading out of the development time distribution that occurs at high larval density in the JB, and greatly increases mean development time, is likely to occur to a much reduced degree in the FEJ because the transition to wandering stage is time-cued in the selected populations, rather than food/size-cued, as it is in typical *D. melanogaster* popualtions, including the JB controls. We stress this point because it illustrates how increased canalization against variation in a macroenvironmental variable can result as a by-product of selection in a constant environment, rather than being selected for directly. Here, the increased canalization of development time with regard to high larval density in FEJ is due to the fact that FEJ larvae stop feeding when still small, rather than being due to any specific developmental buffering mechanism.

It is important to note that the effect of temperature on individual variation for development time, and how the degree of variation changed from one thermal environment to another, varied widely across replicate populations. Thus, neither did the thermal plasticity of development time evolve in FEJs compared to JBs, nor was the extent of trait variation across temperature consistent from one replicate population to another within a selection regime. All of these indicate an overall lack of canalization for development time against temperature changes in our study populations. This is in contrast with density, where changes in trait variation across treatments were consistent across blocks, and also, FEJ mean development time is more canalized against high density than JBs.

The macroenvironmental effects of rearing temperature and density on mean egg to adult survivorship, a correlated response to selection, in the FEJ and JB reveal a pattern different from that seen for egg to adult development time, the direct reponse to FEJ selection (Figure 4). Unlike for development time, variation in egg to adult survivorship across densities is greater than that across temperatures (Figure 6). Moreover, there is no evidence for greater canalization of egg to adult survivorship in FEJ or JB with regard to macroenvironmental variation in either temperature or density.

Turning now to the sensitivity of FEJ and JB development time to microenvironmental variation across vials, we find no evidence that the microenvironmental sensitivity of development time differs between selection regimes or temperature and density levels (Figure 2; Tables 2,3). The sensitivity of FEJ and JB egg to adult survivorship is much higher and also more variable across selection regimes, temperatures and densities (CV 4-15% *vs*. 1% for development time), although only the effects of selection regime and the selection by density interaction are significant (Figure 5; Tables 6,7). Mean survivorship tends to reduce at 300 eggs per vial to a similar degree in both the JB and FEJ, compared to the lower densities (Figure 4), and so does the across vial CV in survivorship, but only in the FEJ (Figure 5). Interestingly, the CV of survivorship for FEJ is typically higher than that for JB in all combinations of temperature and density, except at 300 eggs per vial at 18 and 25°C (Figures 5 and 6), suggesting that the microenvironmental sensitivity of this trait in FEJ is greater than in the ancestral JB controls. This is possibly because egg to adult survivorship is very strongly correlated with fitness in the JB, in which all adults eclosing contribute to the next generation as opposed to just the first few percent eclosing in the FEJ. Consequently, in the FEJ, the primary fitness correlate is development time, not egg to adult survivorship: a variant with really fast development and a small survival cost will likely be favoured by selection in the FEJ regime. The cause for FEJ CV in survivorship being less than JB at 300 eggs per vial could be a result of that density being more stressful for JB than FEJ, due to the reduced size and enhanced carrying capacity of the latter (see above). Thus, the increased sensitivity of JB survivorship at high density, compared to FEJ, to microenvironmental variation across vials is likely to be specifically due to JB being more susceptible to mortality at 300 eggs per vial than FEJ.

Finally, when we examine the effects of selection regime, temperature and density on among individual within vial variability of development time in this study, the main finding is that the temperature effect on CV, unlike that on the mean development time (Figure 1) is less than that of density, and similar on FEJ and JB (Figure 3). Moreover, the sensitivity of among individual variation to increasing density is substantially greater in JB than in FEJ, indicating that FEJ have evolved reduced ultra-microenvironmental canalization of development time with regard to larval density, in tandem with increased macroenvironmental canalization. Again, this is likely a reflection of the much greater effect of high density on JB larvae, due to their larger size and food/size-cued transition from feeding to wandering, as contrasted to the small size of FEJ larvae with a time-cued feeding to wandering transition.

In the FEJ the nature of selection for development time was directional, at least for the first few hundred generations, although the selection response did eventually plateau (A. Joshi, *pers*. *obs*.). While stabilizing selection has been predicted to result in canalization of a trait, directional selection is often thought to lead to decanalization (Schmalhausen, 1949; Rice, 1998; Pélabon *et al.*, 2010; Hayden *et al.*, 2012). Our results, however, suggest that even directional selection can lead to canalization of a trait, as the FEJ appear to have evolved canalization of development time with regard to larval density, both in terms of mean trait value and among individual variability in trait value. More importantly, increased canalization of development time with regard to larval density in FEJ, relative to the ancestral JB controls, seems to be due to the specific effects of the restructuring of the pre-adult life-stage durations and reduction of body size at pupariation. Moreover, the magnitude of effects of temperature and density on mean development time and CV of development time across individuals were opposite. Overall, then, our results underscore that whether or not canalization results from directional selection may often depend on the trait in question, the environmental factor under study, the definition of canalization, and specific details of the ecology and physiology of the populations studied. Some other recent findings also suggest that directional and disruptive selection may not always lead to decanalization as predicted by theory (Hansen *et al.*, 2006; Hayden *et al.*, 2014). Some caution is, therefore, in order when drawing generalized conclusions about the relationship between different forms of selection and canalization from a hypothetical perspective:such relationships be need to be explored on a case-by-case basis and canalization, like evolution in general, may turn out to be ‘local’ (sensu Rose *et al*., 2005). Overall the understanding of the mechanistic basis of canalization remains extremely poor, and to add to that, the link between selection and canalization is equally obscure (Stearns, 2002). We need lot more theoretical and empirical work investigating both mechanisms and causes of canalization, and its evolution, to have a clearer understanding of the phenomenon.

In our study, we focused on classic life-history traits that are fitness correlates. Historically, though, studies of canalization havetypically focused on morphological traits. Waddington (1953, 1956) pioneered the concept of canalization and demonstrated the phenomena of canalization through his experiments on the morphological phenotypes ‘crossveinless’ and ‘bithorax’ in *Drosophila* (Waddington, 1953, 1956). Thus, the concept of canalization was originally proposed and subsequently studied to describe the robustness of developmental pathways governing morphological characters (Dunn and Fraser, 1958; Rendel, 1959). However, the basic criterion for canalization is robustness of phenotypic expression or maintenance of low phenotypic variation of a trait in the face of mutational perturbation or environmental variation. Given such criteria, canalization in principle should be traceable for any quantitative trait, whether or not morphological. However, studies addressing canalization of morphological traits are more common compared to other traits (Dunn and Fraser, 1958; Rendel, 1959; Gibson and Hogness, 1996; Dworkin, 2005a, b; Hall *et al.*, 2007; Sgró *et al.*, 2010; Takahashi *et al.*, 2010; Groth *et al.* 2018). Besides, demonstration of canalizing effect of the chaperone protein Hsp90 in *Drosophila* (Rutherford and Lindquist, 1998) and *Arabidopsis* (Sangster *et al.*, 2008) on a wide variety of morphological traits provided the first evidence of canalizing mechanisms at the molecular level. Recent literature shows an upsurge in interest on canalization in molecular biology as canalization is being investigated for suite of traits ranging from transcript abundance to gene regulatory networks to ribozyme diversity (Hayden *et al.* 2014; Félix and Barkoulas, 2015). However, studies exploring canalization of complex life-history traits are rare. Stearns and Kawecki (1994) and Stearns *et al.* (1995) investigated the canalization of life-history traits in *Drosophila* over two decades ago. In the past decade, few studies addressed canalization of complex traits like reproductive output, body size, developmental period and mate preference (Baer, 2008; Takahashi *et al.*, 2011; Svensson *et al.* 2014). Canalization, thus, needs to be viewed as a general phenomenon controlling trait variation, connecting genotypes to phenotypes, and limiting studies of canalization to morphological traits alone may hinder achieving a broader understanding of the phenomenon, especially its relationship to different forms of selection.

## Conclusion

Our study is exploratory in nature and it sheds light on a number of aspects of the relationship between selection, canalization and plasticity. We demonstrate that (a) a quantitative trait (*e.g.* development time) can evolve without any change in its plasticity (*e.g.* thermal plasticity), (b) canalization can evolve for a life-history trait (*e.g.* development time) in laboratory populations within a span of few hundred generations, (c) contrary to expectations, directional selection can lead to environmental canalization (against developmental density or crowding) of a life history trait, as a by product of the evolved changes in the life history and physiology of the population, and, (d) a trait can be canalized for one environmental factor (density) but may not be canalized for another (temperature). The last two points show that canalization, like many other biological phenomena could be context specific, and one needs to be cautious before drawing generalized conclusions about the causes and manifestations of canalization. Canalization needs to be viewed as a phenomenon controlling trait variation across different hierarchical levels in biology, starting from cellular molecules to complex traits. However, both proximate and ultimate causes of canalization remain poorly understood, and a concerted effort of theoretical and empirical studies are needed to understand this fascinating phenomenon. And last but not the least, this study also demonstrates experimental evolution can be a powerful tool to understand intricacies of the evolutionary process, with particular focus on trait variation.

## Acknowledgements

We thank Archana Mohan for handling data, Ananda T. for help in the experiment and N. Rajanna and Manjesh for general help in the laboratory. SG was supported by a DST WOS A fellowship (SR/WOS-A/LS-1179/2015(G)) during the preparation of the manuscipt, and by a fellowship from JNCASR during the experimental work..KMS thanks Council of Scientific and Industrial Research for a senior research fellowship. The experimental work was supported in parts by funds from the Department of Science and Technology, Government of India, and the preparation of the manuscript by a J C Bose National Fellowship of the Science and Engineering Research Board, Government of India, to AJ. We thank three anonymous reviewers for comments on earlier drafts of this manuscript.

## Author contributions statement

SG and AJ conceived of the experiments, SG, KMS, JM carried out the experiments, SG did the data analysis, SG wrote the manuscript, and both SG and AJ carried out subsequent editing of the manuscript.

## References

1. Félix, M., and Barkoulas, M. (2015). Pervasive robustness in biological systems. Nat. Rev. Gen. 16, 483–496.

2. Waddington, C. H. (1942). Canalization of development and the inheritance of acquired characters. Nature 150, 563–565

3. Waddington, C. H. (1961). Genetic assimilation. Adv. Genet. 10, 257–293.

4. Dworkin, I. (2005a). A study of canalization and developmental stability in the sternopleural bristle system of Drosophila melanogaster. Evolution 59, 1500–1509

5. Waddington, C. H. (1956) Genetic assimilation of the bithorax phenotype. Evolution 10, 1–13

6. Falconer, D. S. and Mackay, T. F. (1996) Introduction to Quantitave Genetics. New York: The Ronald Press Company

7. Debat, V., and David, P. (2001). Mapping phenotypes: Canalization, plasticity and developmental stability. TrendsEcol. Evo. 16, 555–561

8. Hall, M. C., Dworkin, I., Ungerer, M. C., and Purugganan, M. (2007). Genetics of microenvironmental canalization in Arabidopsis thaliana. Proc. Natl. Acad. Sci. USA. 104 (34), 13717–13722

9. Morgante, F., Sorensen, P., Sorensen, D. A., Maltecca, C. and Mackay, and T. F. (2015). Genetic architecture of micro-environmental plasticity in Drosophila melanogaster. Sci. Rep. 5, 09785

10. Stearns, S. C., and Kawecki, T. J. (1994). Fitness sensitivity and the canalization of life-history traits. Evolution 48, 1438–1450

11. Wagner, G. P., Booth, G. and Bagheri-Chaichian, H. (1997), A Population Genetic Theory Of Canalization. Evolution 51, 329–347

12. Rutherford, S. L., and Lindquist, S. (1998). Hsp90 as a capacitor for morphological evolution. Nature 396, 336–342

13. Sgrò, C. M., Wegener, B., and Hoffmann, A. A. (2010). A naturally occurring variant of Hsp90 that is associated with decanalization. Proc. Biol. Sci. 277(1690), 2049–2057

14. Waddington, C. H. (1952). Selection for the basis of an acquired character. Nature 169, 278

15. Waddington, C. H. (1953). Genetic assimilation of an acquired character. Evolution 7, 118–126

16. Bateman, K. G. (1959) The genetic assimilation of four venation phenocopies. J. Genet. 56, 443–474

17. Rendel, J. M. (1959). Canalization of the scute phenotype in Drosophila. Evolution 13, 425–439

18. Gibson, G., and Dworkin, I. (2004). Uncovering cryptic genetic variation. Nat. Rev. Genet. 5, 681–690

19. Woltereck, R. (1909). Weitere experimentelle Untersuchungen űber Artveränderung, speziell űberdas Wesen quantitativer Artunterschiede bei Daphniden. Verh. D. Tsch. Zool. Ges. 19, 110–172

20. Proulx, S. R., and Phillips, P. C. (2005). The opportunity for canalization and the evolution of genetic networks. Am. Nat. 165, 147–162

21. Dor, D., and Jablonka, E. (2010). Plasticity and canalization in the evolution of linguistic communication: An evolutionary developmental approach. In R. Larson, V. Déprez, & H. Yamakido (Eds.), The Evolution of Human Language: Biolinguistic Perspectives (Approaches to the Evolution of Language, pp. 135–147). Cambridge: Cambridge University Press

22. McDonald, J. M., Ghosh, S. M., Gascoigne, S. J., and Shingleton, A. W. (2018). Plasticity through canalization: The contrasting effect of temperature on trait size and growth in Drosophila. Front. Cell Dev. Biol. 6(156)

23. de Visser, J. A. G. M., Hermisson, J., Wagner, G. P., Ancel Meyers, L., Bagheri-Chaichian, H., Blanchard, J. L., Chao, L., Cheverud, J. M., Elena, S. F., Fontana, W. et al. (2003). Perspective: Evolution and detection of genetic robustness. Evolution 57, 1959–1972

24. Flatt, T. (2005). The evolutionary genetics of canalization. Q. Rev. Biol. 80, 287–316

25. West-Eberhard, M. J. (2003). Developmental Plasticity and Evolution. Oxford University Press, New York

26. Pigliucci, M. (2005). Evolution of phenotypic plasticity: where are we going now? Trends Ecol. Evol. 20, 481–486

27. Fusco, G., and Minelli, A. (2010). Phenotypic plasticity in development and evolution: facts and concepts. Phil. Tran. R. Soc. B 365(1540), 547–556

28. Moczek, A. P., Sultan, S., Foster, S., Ledon-Rettig, C., Dworkin, I., Nijhout, H. F,. Abouheif, E., and Pfennig, D. W. (2011). The role of developmental plasticity in evolutionary innovation. Proc Biol. Sci. 278(1719), 2705–2713

29. Paaby A. B., and Testa N. D. (2018) Developmental Plasticity and Evolution. In: Nuno de la Rosa L., Müller G. (eds). Evolutionary Developmental Biology. Springer, Cham, 1–14

30. Debat, V. and Le Rouzic, A. (2018). Canalization, a central concept in biology. Sem. Cell. Dev. Biol. 08–22

31. Groth, B. R., Huang, Y., Monette, M. J., and Pool, J. E. (2018). Directional selection reduces developmental canalization against genetic and environmental perturbations in Drosophila wings. Evolution 72(8), 1708–1715

32. Fossen, E. I., Pélabon, C. and Einum, S. (2018). An empirical test for a zone of canalization in thermal reaction norms. J. Evol. Biol., 31, 936–943

33. Jiang P., Kreitman, M., and Reinitz, J. (2018). The relationship between robustness and evolution. bioRxiv 268862; doi: 10.1101/268862

34. Eshel, I., and Matessi, C. (1998). Canalization, genetic assimilation and preadaptation: a quantitative genetic model. Genetics 149, 2119–2133

35. Siegal, M. L. and Bergman, A. (2002). Waddington’s canalization revisited: developmental stability and evolution. Proc. Natl. Acad. Sci. USA 99, 10528–10532

36. Rendel, J. M. (1967). Canalization and gene control. New York: Logos Press

37. Schmalhausen I. I., (1949). Factors of Evolution: The Theory of Stabilizing Selection, Univ. of Chicago Press, Chicago, reprinted (1986)

38. Dworkin, I. (2005b). Evidence for canalization of distal-less function in the leg of Drosphilia melanogaster. Evol. Dev. 7, 89–100

39. Dworkin, I. (2005c). Canalization, cryptic variation and developmental buffering: a critical examination and analytical perspective. (B. Hallgrimsson and B. K. Hall, eds.. Variation. Elsevier

40. Shaw, J. R., Hampton, T. H., King, B. L., Whitehead, A., Galvez, F., Gross, R. H., Keith, N., Notch, E., Jung, D., Glaholt, S. P., Chen, C. Y., Colbourne, J. K., and Stanton, B. A. (2014). Natural Selection Canalizes Expression Variation of Environmentally Induced Plasticity-Enabling Genes. Mol. Biol. Evol. 31(11), 3002–3015

41. Prasad, N. G., and Joshi, A. (2003). What have two decades of laboratory life-history evolution studies on Drosophila melanogaster taught us? J. Genet. 82, 45–76

42. Garland T. Jr., and Rose, M. R. (Eds.) (2009). Experimental Evolution: Concepts, Methods and Applications of Selection Experiments. University of California Press

43. Stearns, S. C., Kaiser, M. and Kawecki, T. J. (1995). The differential canalization of fitness components against environmental perturbations in Drosophila melanogaster. J. Evol. Biol. 8, 539–557

44. Kawecki, T. J., Lenski, R. E., Ebert, D., Hollis, B., Olivieri, I., and Whitlock, M. C. (2012). Experimental Evolution. Trends Ecol. Evol. 27, 547–560

45. Prasad, N. G., Shakarad, M., Gohil, V. M., Sheeba, V., Rajamani, M., and Joshi, A. (2000). Evolution of reduced pre-adult viability and larval growth rate in laboratory populations of Drosophila melanogaster selected for shorter development time. Genet. Res. 76, 249–259

46. Prasad, N. G., Shakarad, M., Anitha, D., Rajamani, M., and Joshi, A. (2001). Correlated responses to selection for faster development and early reproduction in Drosophila: the evolution of larval traits. Evolution 55, 1363–1372

47. Joshi, A., Prasad, N. G., and Shakarad, M. (2001). K-selection, α-selection, effectiveness, and tolerance in competition: density-dependent selection revisited. J. Genet. 80, 63–75

48. Shakarad, M., Prasad, N. G., Gokhale, K., Gadagkar, V., Rajamani, M., and Joshi. A. (2005). Faster development does not lead to correlated evolution of greater pre–adult competitive ability in Drosophila melanogaster. Biol. Lett. 1, 91–94. doi: 10.1098/2004.0261.

49. Ghosh-Modak S. (2009). Evolution of Life-history Traits, Canalization and Reproductive Isolation in Laboratory Populations of Drosophila melanogaster Selected for Faster Pre-adult Development and Early Reproduction. PhD Thesis, Jawaharlal Nehru Centre for Advanced Scientific Research (JNCASR), Bengaluru, India.

50. Ghosh-Modak, S., Satish, K. M., Mohan, J., Dey, S., Raghavendra, N., Shakarad, M., and Joshi, A. (2009). A possible trade-off between developmental rate and pathogen resistance in Drosophila melanogaster. J. Genet. 88(2), 253–256

51. Ghosh S. M. and Joshi A. (2012) Evolution of reproductive isolation as a by-product of divergent life-history evolution in laboratory populations of Drosophila melanogaster. Ecol. Evol. 2(12), 3214–26

52. Dey, P., Mendiratta, K., Bose, J., and Joshi, A. (2016). Enhancement of larval immune system traits as a correlated response to selection for rapid development in Drosophila melanogaster. J. Genet. 95(3), 719–723

53. Sheeba, V., Madhyastha, N. A. A., and Joshi, A. (1998). Oviposition preference for novel versus normal food resources in laboratory populations of Drosophila melanogaster. J. Biosci. 23, 93–100.

54. Ives, P. T. (1970). Further studies on the South Amherst population of Drosophila melanogaster. Evolution 24, 507–518.

55. Rose, M. R., and Charlesworth, B. (1981). Genetics of life history in Drosophila melanogaster. II. Exploratory selection experiments. Genetics 97(1),187–96.

56. Joshi, A., and Mueller, L. D. (1996). Density-dependent natural selection in Drosophila: trade-offs between larval food acquisition and utilization. Evol. Ecol. 10, 463–474

57. Sokal, R. R., and Rohlf, F. J. (1998). Biometry. 3rd ed. Freeman, New York.

58. Shakarad, M., Prasad, N. G., Rajamani, M. and Joshi A. (2001). Evolution of faster development does not lead to greater fluctuating asymmetry of sternopleural bristle number in Drosophila. J. Genet. 80, 1–7

59. Sang, J. H. (1956). The quantitative nutritional requirements of Drosophila melanogaster. J. Expt. Biol., 33, 45–72

60. Ashburner, M. (1989). Drosophila: a Laboratory Handbook, Cold Spring Harbour Laboratory Press, Cold Spring Harbour, New York

61. Dey, S., Dey, S., Mohan, J., and Joshi, A. (2006). Micro-environmental variations in pre-assay rearing conditions can lead to anomalies in the measurement of life-history traits. J. Genet. 85, 53–56 van der Have T. M., and de Jong G. (1996). Adult Size in Ectotherms: Temperature Effects on Growth and Differentiation. J. Theo. Biol. 183(3), 329–340

62. Angilletta, M. J., Jr., Steury, T. D., and Sears, M. W. (2004) Temperature, growth rateand body size in ectotherms: fitting pieces of a life-history puzzle. Integr. Comp. Biol. 44(6), 498–509

63. Ghosh, S. M., Testa, N. D., and Shingleton, A. W. (2013). Temperature-size rule is mediated by thermal plasticity of critical size in Drosophila melanogaster. Proc. Biol. Sci., 280(1760), 20130174

64. Gebhardt, M. D., and Stearns, S. C. (1988). Reaction norms for developmental time and weight at eclosion in Drosophila mercatorum. J. Evol. Biol. 1, 335–354

65. Gibert, P., Capy, P., Imasheva, A., Moreteau, B., Morin, J. P., Pétavy, G., and David, J. R. (2004). Comparative analysis of morphological traits among Drosophila Melanogaster and D. Simulans: Genetic Variability, Clines and Phenotypic Plasticity. Genetica 120,165–179

66. Borash, D. J., Gibbs, A. G., Joshi, A., and Mueller, L. D. (1998). A genetic polymorphism maintained by natural selection in a temporally varying environment. Am. Nat. 151, 148–156

67. Sarangi, M. (2018). Ecological details mediate different paths to the evolution of larval competitive ability in Drosophila, Ph.D. thesis, Jawaharlal Nehru Centre for Advanced Scientific Research (JNCASR), Bengaluru, India

68. Satish, K. M. (2010). Reverse evolution and gene expression studies on populations of Drosophila melanogaster selected for rapid pre-adult development and early reproduction, Ph.D. thesis, Jawaharlal Nehru Centre for Advanced Scientific Research (JNCASR), Bengaluru, India

69. Rice, S. H. (1998), The Evolution Of Canalization And The Breaking Of Von Baer’s Laws: Modeling The Evolution Of Development With Epistasis. Evolution 52: 647–656

70. Pélabon, C., Hansen, T. F., Carter, A. J., and Houle, D. (2010), Evolution Of Variation And Variability Under Fluctuating, Stabilizing, And Disruptive Selection. Evolution 64, 1912–1925

71. Hayden, E.J., Weikert, C., and Wagner, A. (2012). Directional Selection Causes Decanalization in a Group I Ribozyme. PloS One, 7(9), e45351

72. Hansen, T. F., Álvarez-Castro, J. M., Carter, A. J. R., Hermisson, J., and Wagner G. P. (2006). Evolution Of Genetic Architecture Under Directional Selection. Evolution 60(8), 1523–1536.

73. Hayden, E. J., Bratulic, S., Koenig, I., Ferrada, E., and Wagner, A. (2014). The Effects of Stabilizing and Directional Selection on Phenotypic and Genotypic Variation in a Population of RNA Enzymes. J. Mol. Evol. 78, 101–108

74. Rose, M. R., Passananti, H. B., Chippindale, A. K., Phelan, J. P., Matos, M., Teotónio H., and Mueller L. D. (2005). The effects of evolution are local: evidence from experimental evolution in Drosophila. Integr. Comp. Biol. 45, 486–491 Stearns, S. C. (2002). Progress on canalization. Proc. Natl. sci. Acad. USA 99(16), 10229–10230

75. Dunn, R. B., and Fraser A. S. (1958). Selection for an invariant character - 'vibrissae number' - in the house mouse. Nature 181, 1018–1019

76. Gibson, G., and Hogness, D. S. (1996). Effect of polymorphism in the Drosophila regulatory gene Ultrabithorax on homeotic stability. Science 271(5246), 200–3

77. Baer, C. F. (2008). Quantifying the Decanalizing Effects of Spontaneous Mutations in Rhabditid Nematodes. Am. Nat. 172(2), 272–281

78. Takahashi, K. H., Okada, Y., and Teramura, K. (2011) Genome-wide deficiency mapping of the regions responsible for temporal canalization of the developmental processes of. Drosophila melanogaster. J. Hered. 102 (4), 448–57

79. Sangster, T. A., Salathia, N., Lee, H. N., Watanabe, E., Schellenberg, K., Morneau, K., Wang, H., Undurraga, S., Queitsch, C., … Lindquist, S. (2008). HSP90-buffered genetic variation is common in Arabidopsis thaliana. Proc. Natl. Sci. Acad. USA, 105(8), 2969–74

80. Svensson, E. I., Runemark, A., Verzijden, M. N. and Wellenreuther, M. (2014). Sex differences in developmental plasticity and canalization shape population divergence in mate preferences. Proc. Biol. Sci., 281(1797), 20141636–20141636

